# Protein mishandling and impaired lysosomal proteolysis generated through calcium dysregulation in Alzheimer’s disease

**DOI:** 10.1101/2022.08.24.505157

**Authors:** Sarah Mustaly-Kalimi, Robert A. Marr, Alice Gilman-Sachs, Daniel A. Peterson, Israel Sekler, Grace E. Stutzmann

## Abstract

Impairments in neural lysosomal- and autophagic-mediated degradation of cellular debris contribute to neuritic dystrophy and synaptic loss. While these are well-characterized features of neurodegenerative disorders such as Alzheimer’s disease (AD), the upstream cellular processes driving deficits in pathogenic protein mishandling are less understood. Using a series of fluorescent biosensors and optical imaging in model cells, AD mouse models and human neurons derived from AD patients, we reveal a novel cellular signaling cascade underlying protein mishandling mediated by intracellular calcium dysregulation, an early component of AD pathogenesis. Increased Ca^2+^ release via the endoplasmic reticulum (ER) resident ryanodine receptor (RyR) is associated with reduced expression of the lysosome proton pump vATPase subunits (V1B2 and V0a1), resulting in lysosome deacidification and disrupted proteolytic activity in AD mouse models and human induced neurons (HiN). As a result of impaired lysosome digestive capacity, mature autophagosomes with hyperphosphorylated tau accumulated in AD murine neurons and AD HiN, exacerbating proteinopathy. Normalizing AD-associated aberrant RyR-Ca^2+^ signaling with the negative allosteric modulator, dantrolene (Ryanodex), restored vATPase levels, lysosomal acidification and proteolytic activity, and autophagic clearance of intracellular protein aggregates in AD neurons. These results highlight that prior to overt AD histopathology or cognitive deficits, aberrant upstream Ca^2+^ signaling disrupts lysosomal acidification and contributes to pathological accumulation of intracellular protein aggregates. Importantly, this is demonstrated in animal models of AD, and in human iPSC-derived neurons from AD patients. Furthermore, pharmacological suppression of RyR-Ca^2+^ release rescued proteolytic function, revealing a target for therapeutic intervention that has demonstrated effects in clinically-relevant assays.

**Significance Statement:** We demonstrate in model cells, murine neuronal cultures, and iPSC-derived human neurons, that AD associated RyR-Ca^2+^ dyshomeostasis impairs lysosomal acidification, lysosomal proteolytic activity and hinders autophagic-mediated protein aggregate clearance, which are processes vital to neuronal survival. These deficits were reversed by restoring intracellular Ca^2+^ homeostasis. Notably, this provides a therapeutic target and emphasizes the pathogenic relationship between ER-Ca^2+^ handling, that is known to be altered in AD, to pathogenic protein accumulation as a critical turning point in early stages of Alzheimer’s disease.

## Introduction

Aberrant protein aggregation is a common feature in Alzheimer’s disease (AD), where accumulation of abnormal peptide products and damaged organelles further neuropathology (1). Effective clearance of toxic cellular debris is regulated by macroautophagy (henceforth known as autophagy), a major intracellular catabolic pathway in which debris engulfed by autophagosomes is subsequently absorbed and digested by acidic lysosomes via pH-dependent degradative enzymes such as cathepsins, proteases, and hydrolases (2–5). Deficits in the various steps of autophagic clearance, including lysosomal acidification, autophagosome trafficking and biogenesis contribute to aggregation of AD pathogenic protein deposits (6–10). However, the underlying mechanism driving lysosomal dysfunction and protein mishandling remains poorly understood.

In non-dividing post-mitotic cells, such as neurons, autophagy is the primary means of clearing intracellular waste and is thus essential for neuronal viability. Optimal proteolysis is dependent on the acidic lumen (pH ~4.5) of the lysosome which is maintained through an active vacuolar [H+] ATPase (vATPase) complex. (11–13). The neuronal vATPase proton pump is composed of two subunits, including the cytoplasmic V1 and membrane-embedded V0 domains. The V0a1 sub domain is prevalent in the brain (12, 14, 15) and its proper folding within the ER and subsequent trafficking and assembly in the lysosome is regulated by the ER-localized presenilin 1 (PS1) holoprotein (16, 17). This involvement of PS1 is a critical link to AD and points to pathogenic roles beyond its well-characterized function as the enzymatic component of gamma secretase in APP proteolysis. Mutations in PS1 are the most common cause of early-onset familial AD (FAD) (18–20). In addition to generating Aβ42 species, altered PS1 function interferes with V0a1 delivery to the lysosome, resulting in lysosomal alkalization and increased Ca^2+^ release through upregulation of transient receptor potential cation channel mucolipin subfamily member 1 (TRMPL-1). The subsequent activation on calpains and kinases accelerates tau hyperphosphorylation and neurofibrillary tangle (NFT) formation (4, 16, 17, 21–25).

This elevated cytosolic Ca^2+^ resulting from dysfunction of intracellular organelles, such as lysosomes and ER, is a common convergence point in many lysosomal disorders and neurodegenerative disease. Neuronal Ca^2+^ signaling is essential for numerous functions, ranging from gene transcription to apoptosis, and is fundamental to synaptic plasticity, memory encoding, and bioenergetics (26–30). Increased ER-Ca^2+^ release from ryanodine receptors (RyR) is a well-documented mechanism in AD pathogenesis and occurs prior to the emergence of plaques and tau tangles, and contributes to deficits in synaptic structure and function, phospho-tau and Aβ formation, and cognitive decline (31–34). Notably, normalization of RyR-Ca^2+^ signaling in AD models restores synaptic structure and function, reduces protein aggregation and improves cognition and memory performance (32–37).

The physical and functional coupling of the ER to the endolysosomal system allows for Ca^2+^ transfer that regulates vesicular trafficking, nutrient sensing, and autophagy (38–40). Due to the tight communication between ER and lysosomes, and the vast network of both ER and endolysosomes throughout the neuron, alterations in Ca^2+^ signaling can have deleterious consequences on a wide array of cellular functions. Here, we demonstrate a crucial inter-organelle coupling dynamic that reveals how ER-Ca^2+^ release regulates lysosomal proteolysis via maintenance of vATPase activity. Furthermore, this provides a novel mechanism in which upstream ER-Ca^2+^ dysregulation contributes to AD proteinopathy through disruption of this ER-lysosome-autophagosome pathway. This is shown in model cells and AD mouse models, with clinical relevance further demonstrated in iPSC-derived human neurons from AD patients. Lastly, normalization of ER-Ca^2+^ signaling rescues lysosomal proteolysis and autophagosome turnover of hyperphosphorylated tau, thereby identifying an upstream target for AD with broad therapeutic effects.

## Results

Tightly regulated Ca^2+^ signaling is essential for numerous neuronal functions including handling and clearance of misfolded protein aggregates, such as Aβ and hyperphosphorylated-tau (24, 30, 48). In AD, the aberrant Ca^2+^ release from ER stores interferes with lysosomal ionic transport proteins thus, hindering the acidic environment of lysosomes –a crucial component for autophagy-mediated degradation.

### Diminished lysosomal vATPase expression in AD mice is restored after normalizing intracellular Ca^2+^ signaling

The vacuolar-ATPase (vATPase) proton pump is essential for maintaining an acidic environment within the lysosome lumen. To measure the density of vATPase staining in the dorsal hippocampal subfields (CA1, CA3, DG) (49) and cortex, immunohistochemistry was performed to stain for both the membrane-embedded domain (V0a1) and the cytosolic domain (V1B2). An underlying objective is to identify mechanistic deficits early in the disease process, thus, 3-4 month aged-old male and female 3xTg-AD mice and non-transgenic (NTg) controls were used. Our findings demonstrate a significant decrease in hippocampal and cortical V0a1 expression in 3xTg-AD untreated mice relative to NTg untreated mice **(Figure 1A,C;** n=8 animals, CA1 F_(3,26)_=11.23, p<0.0001; CA3 F_(3, 27)_=25.37, p<0.0001,; DG F_(3,24)_=44.08, p<0.0001; Cortex F_(3,27)_=28.34, p<0.0001). In this experiment, and in all assays, we measured sex as a variable, and found no differences between male and female mice. Similarly, there is a significant reduction of V1B2 in 3xTg-AD untreated mice relative to the NTg controls **(Figure 1B,D;** n=8, CA1 F(3,26)=39.06, p<0.0001; CA3 F_(3,26)_=27.57, p<0.0001; DG F_(3,26)_=29.12, p<0.0001; Cortex F_(3,28)_=28.99, p<0.0001). Since these vATPase changes may reflect disruption in lysosomal turnover, a lysosomal surface marker, LAMP1, was additionally immunolabeled. However, there were no significant differences in relative density of lysosomes **(Figure 1E;** n=8, CA1 F_(3,26)_=2.58, p=0.076; CA3 F_(3,24)_=1.797, p=0.17; DG F_(3,22)_=2.34, p=0.10; Cortex F(3,28)=1.70, p=0.19). Thus, independent of lysosomal turnover, vATPase expression was reduced in the dorsal hippocampus and overlying cortex in the AD mice, consistent with disruption in vATPase trafficking and function prior to histopathological deposits.

**Figure 1.**
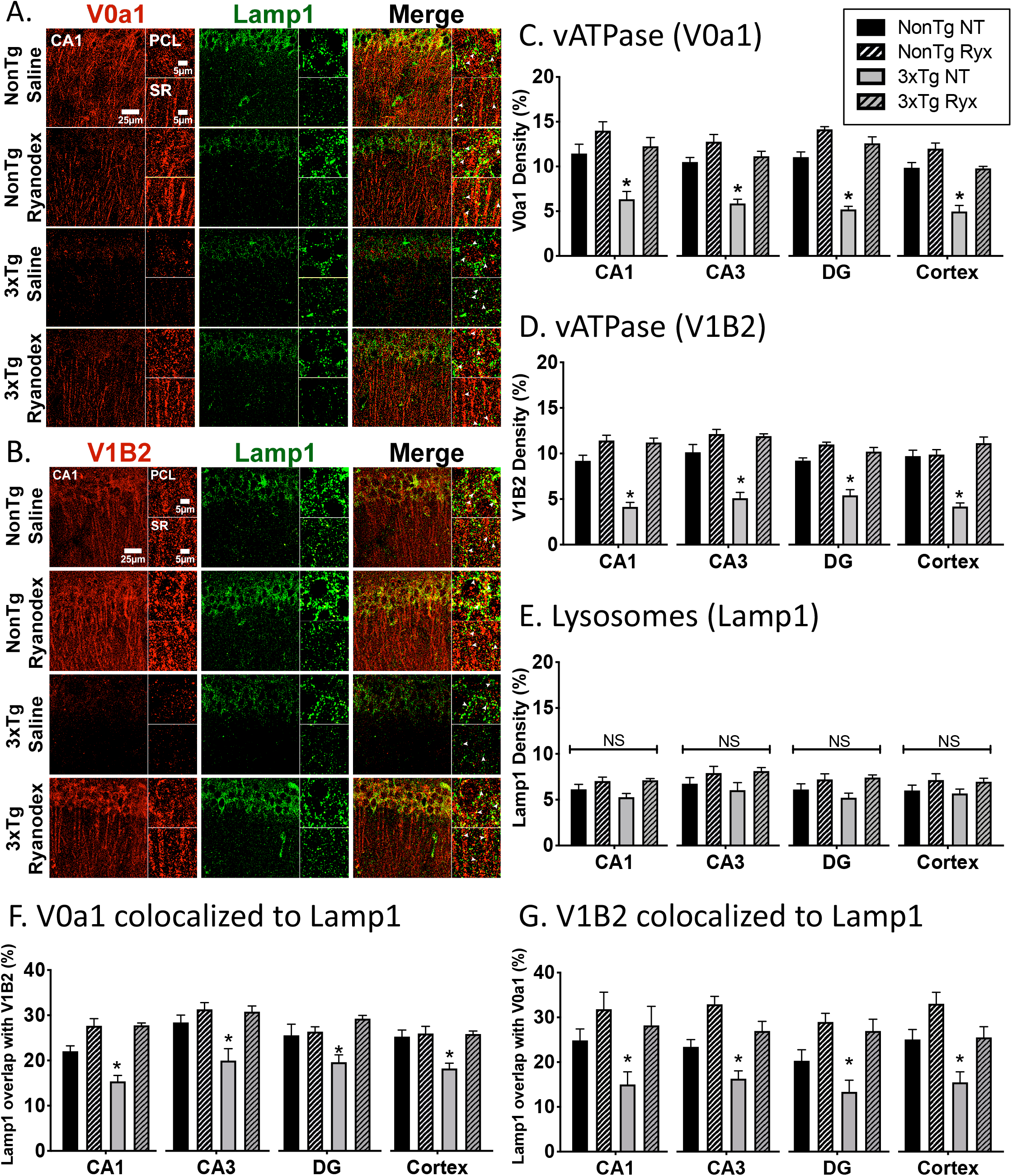
Decreased vacuolar-ATPase in AD mice are restored after normalizing intracellular Ca^2+^ levels. Lysosomal proton pump, vATPase, expression is dependent on regulated RyR-Ca^2+^ signaling. Hippocampus from 3-4-month-old 3xTg-AD mice and age-matched NTg controls were immunolabled for vATPase membrane-embedded V0a1 and cytosolic V1B2 subunits along with lysosomal surface marker (Lamp1). (A) Representative high-resolution images were taken on Leica SP8 63X with 441X zoom of the primary cellular layer (PCL) and stratum radiatum (SR) of the CA1 field. Quantification showed decreased fluorescence density (% of threshold area over background) for (C) V0a1, (D) V1B2 in multiple regions of the hippocampus (CA1, CA3, DG) and cortex; that was rescued with Ryanodex treatment (10mg/kg; 30-day ip). (E) There were no significant differences in Lamp1 density throughout the hippocampus and cortex. Colocalization of (F) V0a1 and (G) V1B2 to lysosomes specifies that deficits seen in vATPase are localized to lysosomes. (NTg Vehicle, black bar, n=8 animals; NTg Ryanodex, black pattern bar, n=8 animals; 3xTg Vehicle, grey bar, n=8 animals; 3xTg Ryanodex, grey pattern bar, n=8 animals) *\*p <0.05*

To confirm the reduced vATPase expression measured localized to lysosomes, as opposed to other acidic compartments such as synaptic vesicles or endosomes, we measured the degree of colocalization between Lamp1 and vATPase subunits, which showed a decreased presence of V0a1 colocalized to lysosomes **(Figure 1F;** n=8, CA1 F(3,22)=4.16, p=0.018; CA3 F_(3,26)_=13.06, p<0.0001; DG F_(3,26)_=7.83, p=0.0007; Cortex F_(3,26)_=8.67, p=0.0004); and V1B2 colocalized to lysosomes **(Figure 1G**, n=8, CA1 F_(3,23)_=22.29, p<0.0001; CA3 F_(3,20)_=8.27, p<0.0009; DG F_(3,22)_=6.73, p=0.002; Cortex F_(3,27)_=8.04, p=0.0006) in all regions of the dorsal hippocampal subfields and cortex in 3xTg-AD untreated mice relative to NTg controls.

Coupling and communication among intracellular organelles is necessary for cellular homeostasis, and likewise, functional impairment in one organelle can impact several cellular systems. It is well-established in AD models that RyR-evoked ER Ca^2+^ release is markedly increased (27, 32, 50–52), and this can be normalized by treatment with negative allosteric RyR modulators such as dantrolene or Ryanodex (32, 35, 37). To determine if the decreased lysosome vATPase expression is a consequence of AD-associated aberrant ER-Ca^2+^ signaling, mice were treated with Ryanodex (10mg/kg; 30-day i.p.) for 4 weeks, and levels of the vATPase subunits were measured and compared between mouse strains. In 3xTg-AD mice, the density of V0a1 **(Figure 1A,C)** and V1B2 **(Figure 1B,D)** is restored to control levels upon normalization of ER Ca^2+^ signaling and similarly, lysosomes colocalized to V0a1 **(Figure 1F)** and V1B2 **(Figure 1G)** are restored to levels comparable to NTg mice. Overall, these data demonstrate that ER-Ca^2+^ dynamics regulate vATPase expression on lysosomes, and aberrant RyR-Ca^2+^ signaling plays a key role in the loss of vATPase and downstream lysosomal function.

### AD-associated aberrant RyR-Ca^2+^ release alkalizes lysosomal pH in model cells and neuronal cultures

Based on the observed deficits in lysosomal vATPase expression in AD models, we further explored the effects of brain-resident RyR isoforms on lysosomal acidity and function. In particular, increased RyR2 expression and changes in secondary structure leading to channel leakiness are associated with AD pathogenesis (37, 53–55). Here, RyR2- and RyR3-expressing HEK cells (42, 43) were assessed for peak RyR-evoked Ca^2+^ release evoked with 10mM caffeine and measured with Fura-2AM. Ryanodex (10μM; 1hr) attenuated the RyR-evoked Ca^2+^ release in both RyR2 **(Figure 2A-B;** n=6, F_(2,15)_=8.63, p=0.0032) and RyR3 cell lines (**data not shown;** n=6, F_(2,15)_=5.80, p=0.014). Inhibiting vATPase activity with bafilomycin treatment (125nM; 4hr) did not significantly alter RyR-evoked Ca^2+^ release in RyR2 or RyR3 lines, confirming that bafilomycin does not have off-target effects on RyR-evoked Ca^2+^ release.

**Figure 2.**
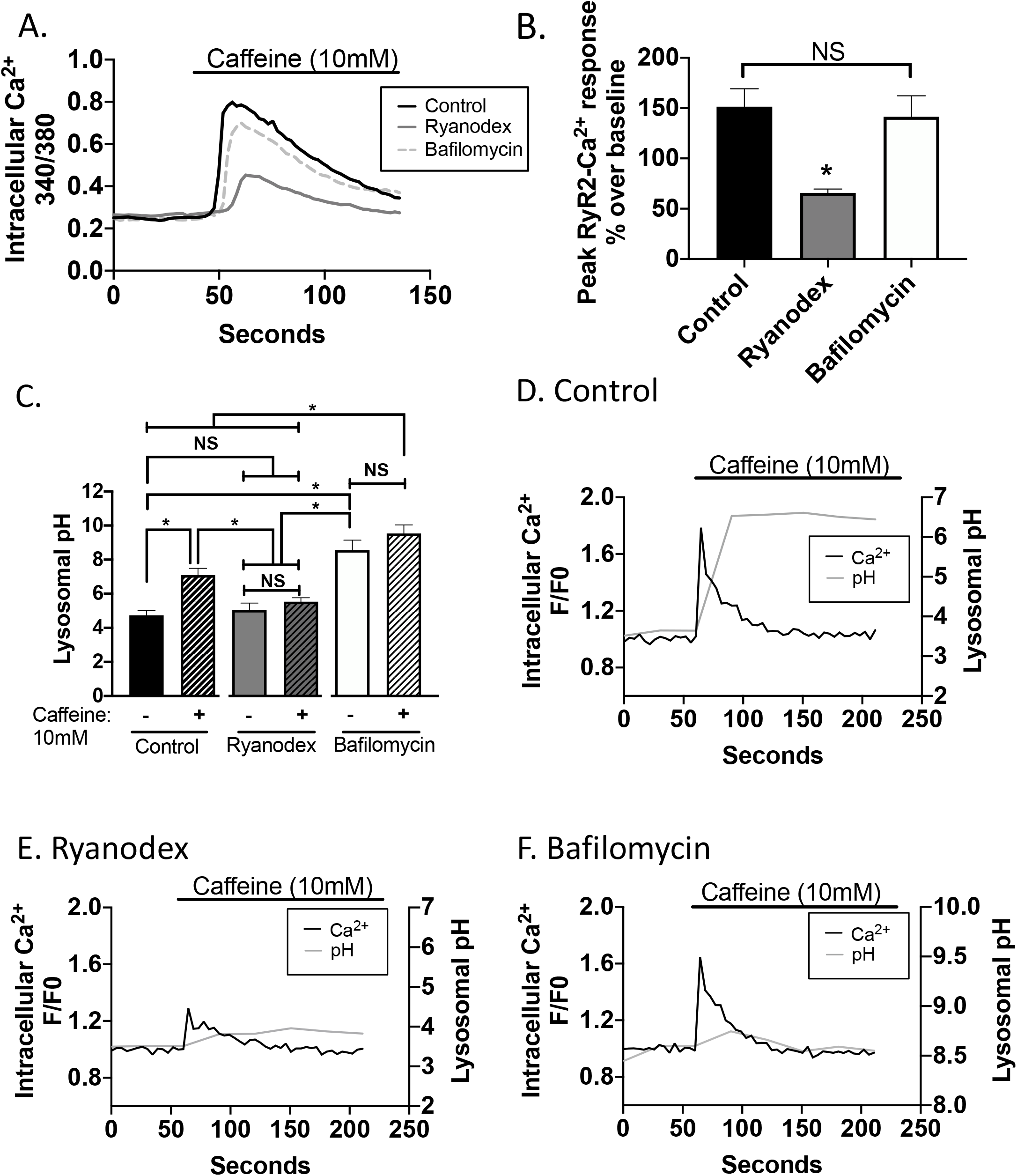
Ryanodine Receptor (RyR)-evoked Ca^2+^ release alkalizes lysosomal pH through vacuolar-ATPase proton pump interference. Modulating RyR-evoked Ca^2+^ release rescues lysosomal alkalization in model cells. (A) Representative traces of RyR-evoked Ca^2+^ release (10mM caffeine) on HEK293T cells overexpressing hippocampal-relevant RyR2 treated with control, Ryanodex or bafilomycin. (B) Cells were incubated with Fura-2AM and ER-Ca^2+^ release was measured as peak change in 340/380 ratio over baseline (n= 6 coverslips/treatment). Normalizing RyR-Ca^2+^ release with Ryanodex treatment (10μM; 1hr) attenuated RyR-evoked Ca^2+^ release. Inhibiting vATPase with bafilomycin (125nM; 4hr) did not affect RyR-evoked Ca^2+^ release. Lysosomal pH was measured using LysoSensor Yellow/Blue dextran (n=27 wells/treatment). (D-F) Representative traces overlaying RyR-Ca^2+^ release and lysosomal pH indicate immediate lysosomal pH fluctuations with a rise in intracellular Ca^2+^ levels. (C,D) RyR-evoked Ca^2+^ release alkalized lysosomal pH. (C,E) Normalizing RyR-Ca^2+^ release with Ryanodex treatment (10μM; 1hr) attenuated the RyR-mediated lysosomal alkalization. (C,F) Inhibiting vATPase with bafilomycin (125nM; 4hr) alkalized lysosomal pH, and no additional pH changes occurred in response to RyR-evoked Ca^2+^ release. *\*p>0.05*

To determine how RyR-Ca^2+^ release impacts lysosomal acidity, lysosomal pH was measured using LysoSensor Yellow/Blue dextran. Lysosomal pH became more alkaline in response to a RyR-evoked Ca^2+^ release in RyR2 **(Figure 2C;** pH of control 4.7±0.28; pH of control + caffeine 7.1± 0.41; n=27, F_(5,156)_ =22.48; p<0.0001) and RyR3 cell lines **(data not shown);** control 3.7±0.35; control + caffeine 6.0± 0.61; n=32, F_(5,187)_=10.27; p<0.0001).

Ryanodex prevented the alkalization of lysosomal pH in both RyR2 (pH with Ryanodex 5.0±0.41; pH with Ryanodex + caffeine 5.5± 0.23) and RyR3 cell lines (pH with Ryanodex 3.5±0.51; pH with Ryanodex + caffeine 3.8± 0.46). The close temporal coupling of these events is demonstrated in **Figure 2D**, in which the evoked Ca^2+^ trace overlaid on the lysosomal pH trace shows the surge of ER-Ca^2+^ release and the resulting rapid alkalization in lysosomal pH. Ryanodex treatment decreases the RyR-Ca^2+^ release and protects from a profound lysosomal alkalization **(Figure 2E)**. Together, these data demonstrate that lysosomal pH is tightly coupled to ER Ca^2+^ signaling and that communication between the ER and lysosome is necessary for lysosomal homeostatic regulation.

As a positive control, inhibiting vATPase with bafilomycin alkalized lysosomal pH, and no additional pH changes occurred in response to RyR-evoked Ca^2+^ release in RyR2 cell lines (**Figure 2C,F)** (pH with bafilomycin 8.5±0.58; pH with bafilomycin + caffeine 9.5±0.51) and RyR3 cell lines (**Supplemental Figure 1C)** (pH with bafilomycin 6.1±0.39; pH with bafilomycin + caffeine 7.2±0.57). This indicates that after blocking proton import via vATPase, subsequent Ca^2+^ release is not acting through alternative ion channels or mechanisms. To establish these lysosomal pH changes are mediated specifically by caffeine-evoked RyR-Ca^2+^ release, doxycycline was not added to the cells thus minimizing RyR overexpression. No significant changes in lysosomal pH after caffeine stimulation were found in HEK293T RyR2 (n=6, t(1,10)=0.036; p=0.97; two-tailed t-test) or RyR3 cells without doxycycline (n=6, t(1,14)=0.006; p=0.10; two-tailed t-test) (data not shown).

Overall, the above data reveal that lysosomal pH is modulated by intracellular ER-Ca^2+^ release. Elevations in RyR-Ca^2+^ release alkalizes lysosomal pH and blocking RyR-Ca^2+^ release diminishes lysosomal alkalization. To validate these findings in a neuronal model, primary neuronal hippocampal cultures were generated from 3xTg-AD and NTg pups. RyR-evoked Ca^2+^ release in these neurons were measured using Fura-2AM and bath application of 10mM caffeine. 3xTg-AD neurons have significantly increased RyR-evoked Ca^2+^ responses compared to NTg neurons **(Figure 3A,B;** NTg control 8.19% ±1.23% over baseline; 3xTg-AD control 25.74%±5.01% over baseline; n=19, F_(5,108)_=8.38; p<0.0001), effects that were mitigated and comparable to NTg controls with Ryanodex treatment (10μM; 1hr) in 3xTg-AD neurons (3xTg-AD Ryanodex 3.01%±0.48% over baseline; NTg Ryanodex 9.65%±2.02% over baseline). Bafilomycin (125nM; 4hr) did not significantly alter RyR-evoked Ca^2+^ release in either 3xTg-AD (28.47%±16.54% over baseline) or NTg (8.62% ±2.99% over baseline) neurons, again showing minimized interference of bafilomycin to RyR-evoked Ca^2+^ release. These results are consistent with previous studies that demonstrated altered upstream ER-Ca^2+^ homeostasis in multiple AD mouse and human neuronal models. This dysregulated ER-Ca^2+^ signaling disrupted synaptic transmission and plasticity, propagated inflammatory responses, and drove pathogenic amyloid and tau formation; all of which was normalized with Ryanodex treatment in the AD models (32, 35, 44, 50, 52).

**Figure 3.**
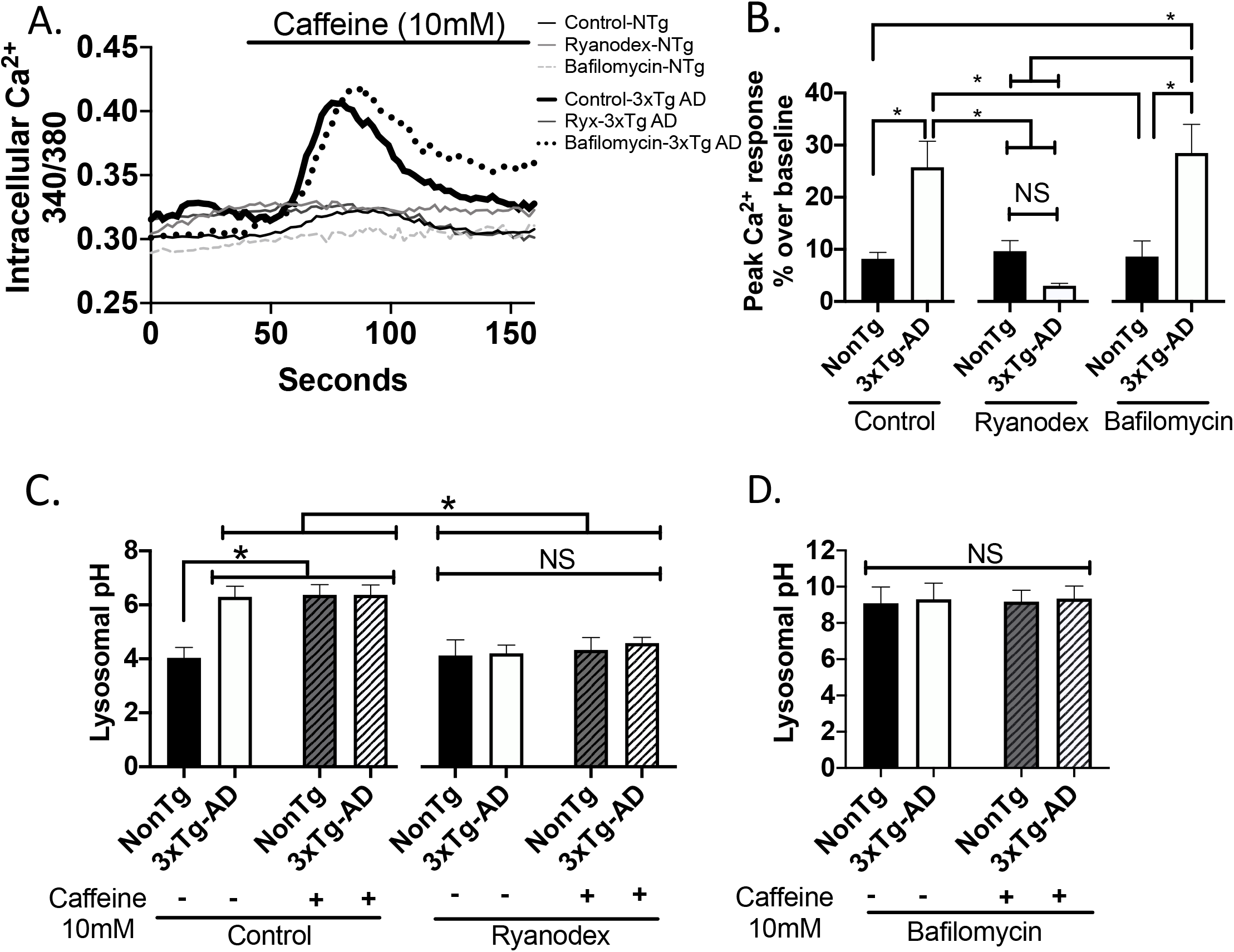
Ryanodine Receptor (RyR2)-evoked Ca^2+^ release alkalizes lysosomal pH through vacuolar-ATPase proton pump interference in 3xTg-AD neuronal cultures. Modulating RyR-Ca^2+^ release regulates lysosomal pH in primary hippocampal murine cultures. (A) Representative traces of RyR-evoked Ca^2+^ release (10mM caffeine) on 3xTg-AD and NTg hippocampal primary neuronal cultures (12-14 days) treated with control, Ryanodex, or bafilomycin. (B) Neurons were incubated with Fura-2AM and ER-Ca^2+^ release was measured as peak change in 340/380 ratio over baseline (n= 19 coverslips/treatment). Normalizing RyR-Ca^2+^ release with Ryanodex treatment (10μM; 1hr) attenuated RyR-evoked Ca^2+^ release. Inhibiting vATPase with bafilomycin (125nM; 4hr) did not affect RyR-evoked Ca^2+^ release. Lysosomal pH was measured using LysoSensor Yellow/Blue dextran on hippocampal primary neuronal culture (n=8 coverslips/treatment). (C) Baseline lysosomal pH is increased in 3xTg-AD neurons as compared to NTg neurons. RyR-evoked Ca^2+^ release alkalized lysosomal pH in NTg neurons, while no further lysosomal pH changes occurred in 3xTg-AD neurons. Ryanodex treatment (10μM; 1hr) attenuated the RyR-mediated lysosomal alkalization and restored baseline lysosomal pH in 3xTg-AD neurons. Bafilomycin (125nM; 4hr) alkalized lysosomal pH, and no additional pH changes occurred in response to RyR-evoked Ca^2+^ release in NTg nor 3xTg-AD neurons. *\*p>0.05*

Here, we demonstrate that lysosomal dysfunction is another consequence of dysregulated ER-Ca^2+^ release. Baseline lysosomal pH was more alkaline in 3xTg-AD neurons compared to NTg neurons (pH of NTg control 4.0±0.38; pH of 3xTg-AD control 6.3±0.39), and RyR-evoked Ca^2+^ release alkalized lysosomal pH in NTg neurons (pH of NTg control+caffeine 6.367±0.38) to levels comparable to 3xTg-AD neurons. Notably, lysosomal pH did not become further alkalized in response to RyR-evoked Ca^2+^ in 3xTg-AD neurons (pH of 3xTg-AD control +caffeine 6.37±0.36; n=8, F_(7,61)_=8.42, p<0.0001), indicating that a maximal threshold for occluding proton flux is at a steady state in the AD neurons. Decreasing aberrant RyR-Ca^2+^ release with Ryanodex (10μM; 1hr), mitigated the effects on pH in NTg neurons (pH of NTg Ryanodex 4.11±0.58; pH of NTg Ryanodex+caffeine 4.33±0.46) as well as rescued the alkalization of lysosomal pH in the 3xTg-AD neurons (pH of 3xTg-AD Ryanodex 4.19±0.31; pH of 3xTg-AD Ryanodex+caffeine 4.58±0.22), demonstrating that lysosomal acidity is in part regulated by the ER through RyR-Ca^2+^ signaling (**Figure 3C**). As expected, inhibiting vATPase activity with bafilomycin treatment (125nM; 4hr) alkalized lysosomal pH in NTg and 3xTg-AD neurons, and subsequently evoking RyR-Ca^2+^ release did not induce a further change in lysosomal pH in either NTg and 3xTg-AD neurons (pH of NTg bafilomycin 9.08±0.89; pH of NTg bafilomycin+caffeine 9.18±0.63; pH of 3xTg-AD bafilomycin 9.31±0.89; pH of 3xTg-AD bafilomycin+caffeine 9.34±0.69, (n=8, F(3,24)=0.02, p=0.10,**Figure 3D)**.

### AD-associated aberrant RyR-Ca^2+^ release alkalizes lysosomal pH and inhibits lysosomal protease activity in human induced neurons

In order to bridge the translational gap between murine and human models of AD, iPSC-derived human induced neurons (HiN) were generated (44) to establish if similar pathophysiological mechanisms were present in neurons derived from AD patients. Immunoassays were performed to measure vATPase density in HiN and, similar to murine samples, a marked decrease in V0a1 (n=9, F_(3,33)_=13.02, p<0.0001, one-way ANOVA**; Figure 4A,C)** and V1B2 (n=9, F_(3,34)_=11.28, p<0.0001; **Figure 4B,D)** densities were observed in HiN derived from AD patients as compared to HiN from control individuals. Normalization of intracellular Ca^2+^ release with Ryanodex restored vATPase expression in AD HiN to levels comparable with non-AD HiN. Additionally, no significant differences in the density of lysosome staining, as measured by live-cell lysosomal dye LysoTracker Red (n=14, F_(3,52)_=2.379, p=0.0802, One-way ANOVA; **Figure 4E)**, was detected between non-AD and AD HiN. Corresponding to the mouse models, ER-Ca^2+^ signaling regulates lysosomal vATPase expression and the loss of vATPase is tied to AD-associated aberrant RyR-Ca^2+^ release.

**Figure 4.**
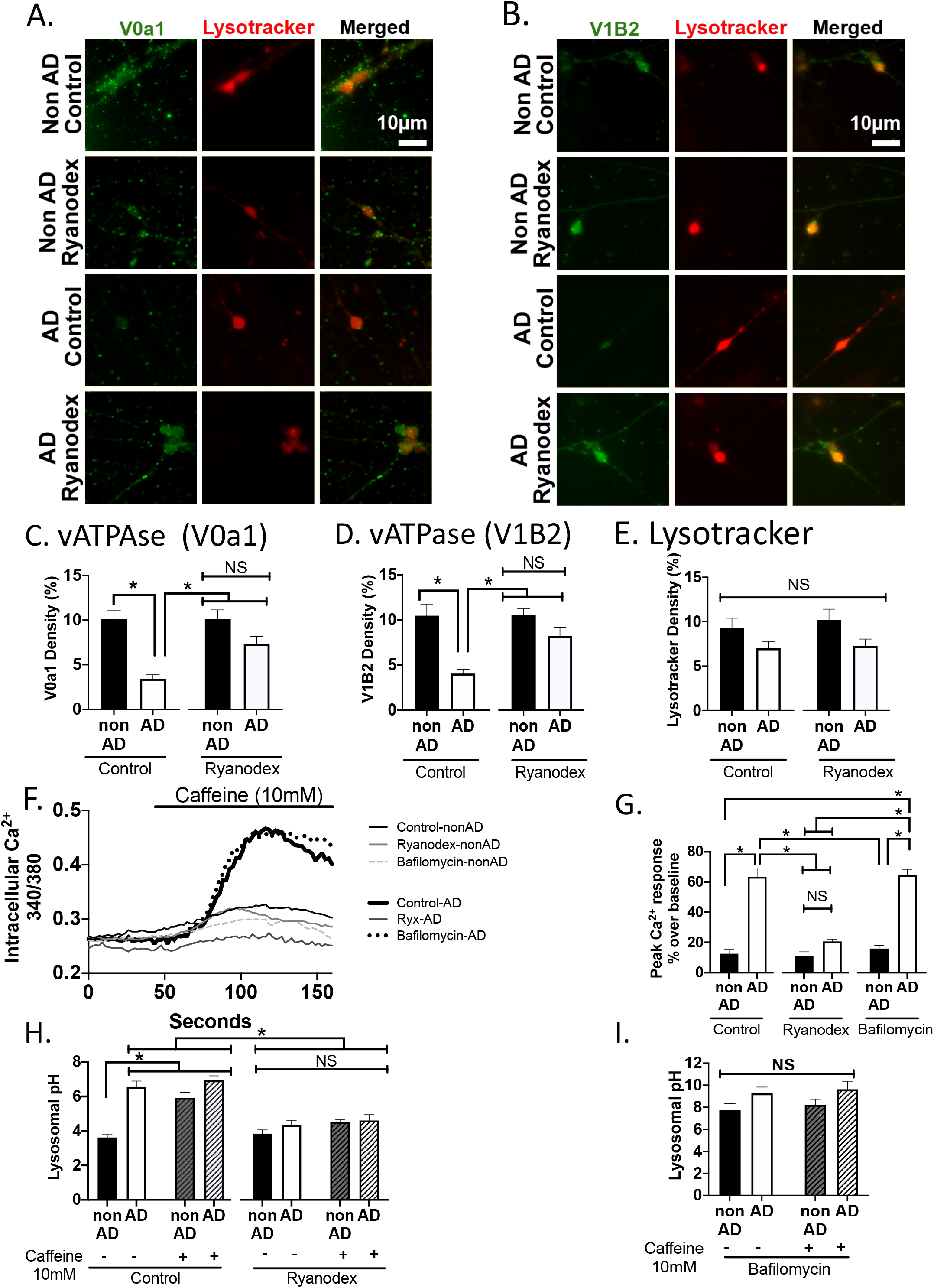
Ryanodine Receptor (RyR)-evoked Ca^2+^ release alkalizes lysosomal pH through vacuolar-ATPase proton pump interference in AD Human Neurons. Modulating RyR-Ca^2+^ regulates vATPase expression and lysosomal pH in human induced neurons (HiN) derived from AD and non-AD patients. HiN were immunolabled for vATPase membrane-embedded V0a1 and cytosolic V1B2 subunits along with a lysosomal marker (Lysotracker). (A) Representative images and quantification showed decreased fluorescence density (% of threshold area over background) for V0a1 (C; n=9 coverslips/treatment), V1B2 (D; n=9 coverslips/treatment), and Lysotracker (E; n=14 coverslips/treatment), that was rescued with Ryanodex (10μM). (F) Representative traces of RyR-evoked Ca^2+^ release (10mM caffeine) on AD and non-AD HiN treated with control, Ryanodex (10μM) or bafilomycin (125nM). (G) Neurons were incubated with ratiometric indicator Fura-2AM and ER-Ca^2+^ release was measured as peak change in 340/380 ratio over baseline (n=8 coverslips/treatment). Normalizing RyR-Ca^2+^ release with Ryanodex treatment (10μM; 1hr) decreased RyR-evoked Ca^2+^ release. Inhibiting vATPase with bafilomycin (125nM; 4hr) did not affect RyR-evoked Ca^2+^ release. Lysosomal pH was measured using LysoSensor Yellow/Blue dextran on HiN (n=11 coverslips/treatment). (C) Baseline lysosomal pH is increased in AD HiN as compared to non-AD HiN. RyR-evoked Ca^2+^ release alkalized lysosomal pH in non-AD HiN, while no further pH changes occurred in AD HiN. Ryanodex treatment (10μM; 1hr) attenuated the RyR-mediated lysosomal alkalization and restored baseline lysosomal pH in AD HiN. Bafilomycin (125nM; 4hr) alkalized lysosomal pH, and no additional pH changes occurred in response to RyR-evoked Ca^2+^ release in non-AD nor AD HiN. *\*p>0.05*

Evoked Ca^2+^ release in HiN was previously characterized (44), and here we show that AD HiN neurons exhibit exaggerated RyR-evoked responses compared to cognitively-normal age-matched HiN controls (**Figure 4F-G**; non-AD HiN 12.53% ±2.74% over baseline; AD HiN control 63.43%±5.86% over baseline; n=8, (F_(5,42)_=55.17; p<0.0001). Ryanodex treatment (10μM; 1hr) restores RyR-evoked Ca^2+^ response in AD HiN (AD HiN Ryanodex 20.64%±1.515% over baseline) to non-AD HiN levels (non-AD HiN Ryanodex 11.17%±2.61% over baseline). Additionally, bafilomycin (125nM; 4hr) did not interfere with RyR-evoked Ca^2+^ release in non-AD nor AD HiN (HiN non-AD bafilomycin 15.87%±2.20% over baseline; HiN AD bafilomycin 64.52%±3.85% over baseline). To examine the role of RyR-Ca^2+^ release on lysosomal pH fluctuation in a clinically-relevant cell system, lysosomal pH was measured using Lysosensor Yellow/Blue dextran. Baseline lysosomal pH in the AD HiN was significantly more alkaline than non-AD HiN (pH of non-AD HiN 3.6±0.17; pH of AD HiN 6.6±0.34). RyR-evoked Ca^2+^ release alkalized lysosomal pH in non-AD HiN, whereas the lysosomal pH did not become more alkaline in AD HiN (pH of non-AD HiN +caffeine 5.9±0.32; pH of AD HiN +caffeine 6.9±0.26), again showing that elevated intracellular Ca^2+^ levels as a result of aberrant RyR activity alkalizes lysosomal pH. Normalizing RyR-Ca^2+^ release with Ryanodex (10μM; 1hr) attenuated the RyR-evoked Ca^2+^ response in non-AD HiN (pH of non-AD HiN Ryanodex 3.84±0.22; pH of non-AD HiN Ryanodex+caffeine 4.52±0.15) as well as rescued the alkalization of lysosomal pH in the AD HiN (pH of HiN AD Ryanodex 4.36±0.262; pH of HiN AD Ryanodex+caffeine 4.61±0.33; **Figure 4H**, n=11, (F_(7,80)_=24.69, p<0.0001). Collectively, this demonstrates that aberrant RyR-Ca^2+^ signaling underlies lysosomal pH deficits in human AD neuronal models, and the AD neurons are, at rest, functioning with compromised lysosomes. As a positive control, bafilomycin alkalized lysosomal pH in both non-AD and AD HiN and RyR-evoked Ca^2+^ release did not further alkalize lysosomal pH in both non-AD and AD HiN **(Figure 4I;** pH of HiN non-AD bafilomycin 7.76±0.56; pH of HiN nonAD bafilomycin+caffeine 8.22±0.49; pH of HiN AD bafilomycin 9.26±0.57; pH of HiN AD bafilomycin+caffeine 9.63±0.73; n=11, (F_(3,40)_=2.22, p=0.10).

Here we demonstrated across various AD models that RyR-Ca^2+^ release alkalizes lysosomal pH, which would be expected to impair lysosomal degradation by inactivating proteases and hydrolases. To confirm, DQ-red BSA was used to measure lysosome protease activity. In AD HiN, protease activity was significantly decreased compared to non-AD HiN. Notably, Ryanodex treatment (10μM; 6hr) rescued protease activity in HiN AD to levels comparable HiN non-AD (**Figure 5 A-B);** n=5, (F_(5,27)_=68.22, p<0.0001, One-way ANOVA). As confirmation, bafilomycin (125nM; 6hr) obliterated protease activity in both non-AD and AD HiN (**Figure 5 A-B**). This, coupled with the above lysosomal pH deficits, supports the hypothesis that AD-associated aberrant RyR-Ca^2+^ signaling disrupts lysosomal function and downstream protein degradation potential in human neurons.

**Figure 5.**
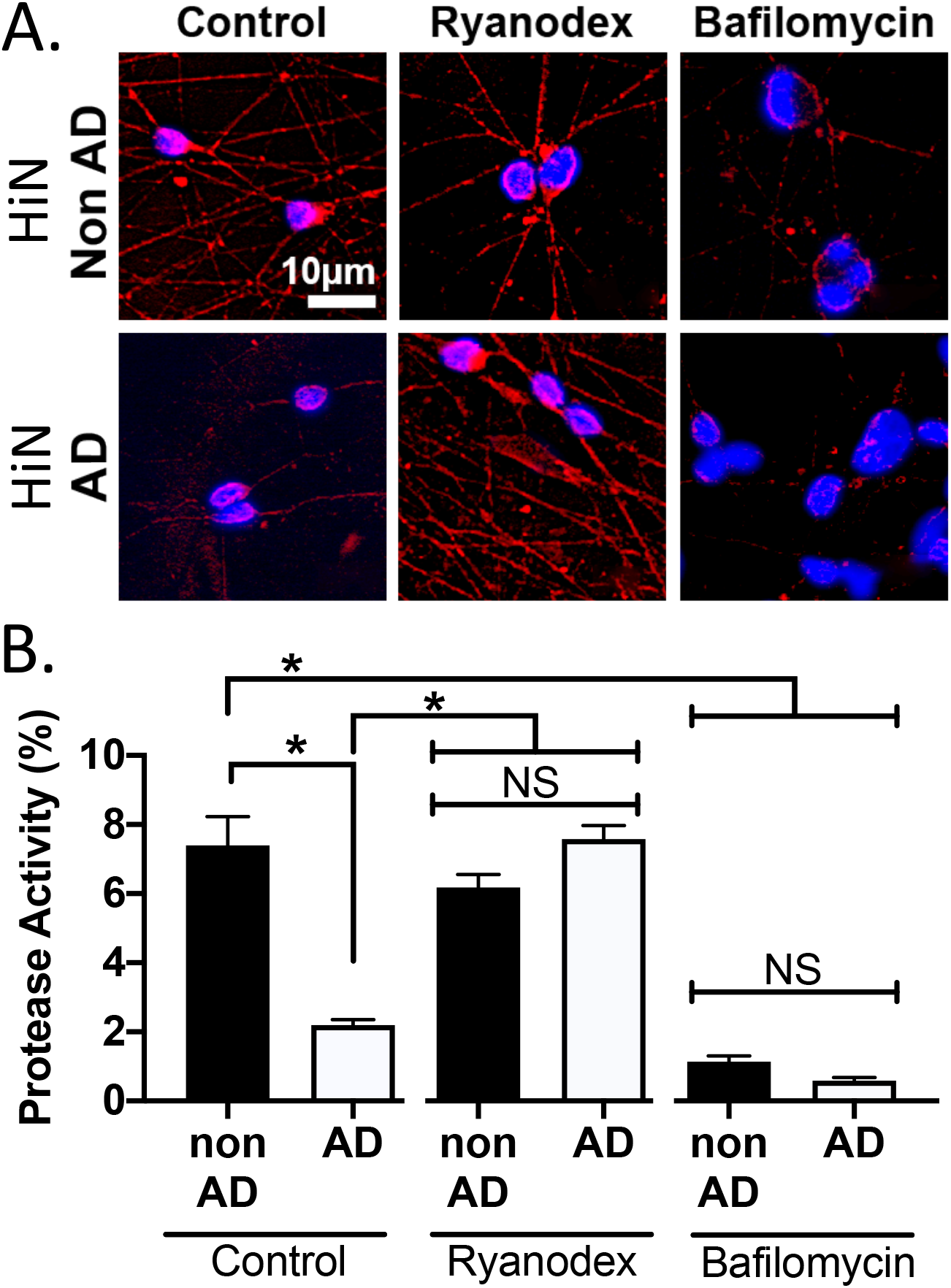
Disrupted lysosomal proteolytic activity in AD human induced neurons. Normalizing RyR-Ca^2+^ release rescues disrupted protease activity in human induced neurons (HiN). (A) Representative images of lysosomal proteolytic activity, measured by DQ-Red BSA, in HiN treated with control, Ryanodex and bafilomycin are shown. (B) Quantification of fluorescence intensity (% signal over background) showed decreased protease activity in AD HiN, which was rescued with Ryanodex (10μM) treatment. vATPase inhibition with bafilomycin (125nM) demolished protease activity in both non-AD and AD HiN. (n=5 coverslips/treatment) *\*p>0.05*.

### Disrupted autophagic-mediated clearance as a result of AD-associated ER-Ca^2+^ release

The alkalization of lysosomes and hindered proteolytic activity impairs the digestion of autophagosomes and their cellular debris. To measure potential deficits in autophagosome clearance, brain slices were immunolabeled for LC3B, a marker for mature autophagosomes, and staining density measured within the dorsal hippocampus (CA1, CA3, DG subfields) and overlying cortex. In 3xTg-AD mice, autophagosome density was significantly increased in all regions examined relative to NTg controls, consistent with a loss of lysosomal proteolytic capacity. Normalizing Ca^2+^ signaling with Ryanodex treatment (10mg/kg, 30-days, i.p.) rescued the density of autophagosomes in 3xTg-AD mice to levels comparable with NTg controls (**Figure 6A-B;** n=8 animals, CA1 (F_(3,28)_=42.76, p<0.0001); CA3 (F_(3, 28)_=47.40, p<0.0001); DG (F_(3,28)_=61.95, p<0.0001); Cortex (F_(3,28)_=54.58, p<0.0001).

**Figure 6.**
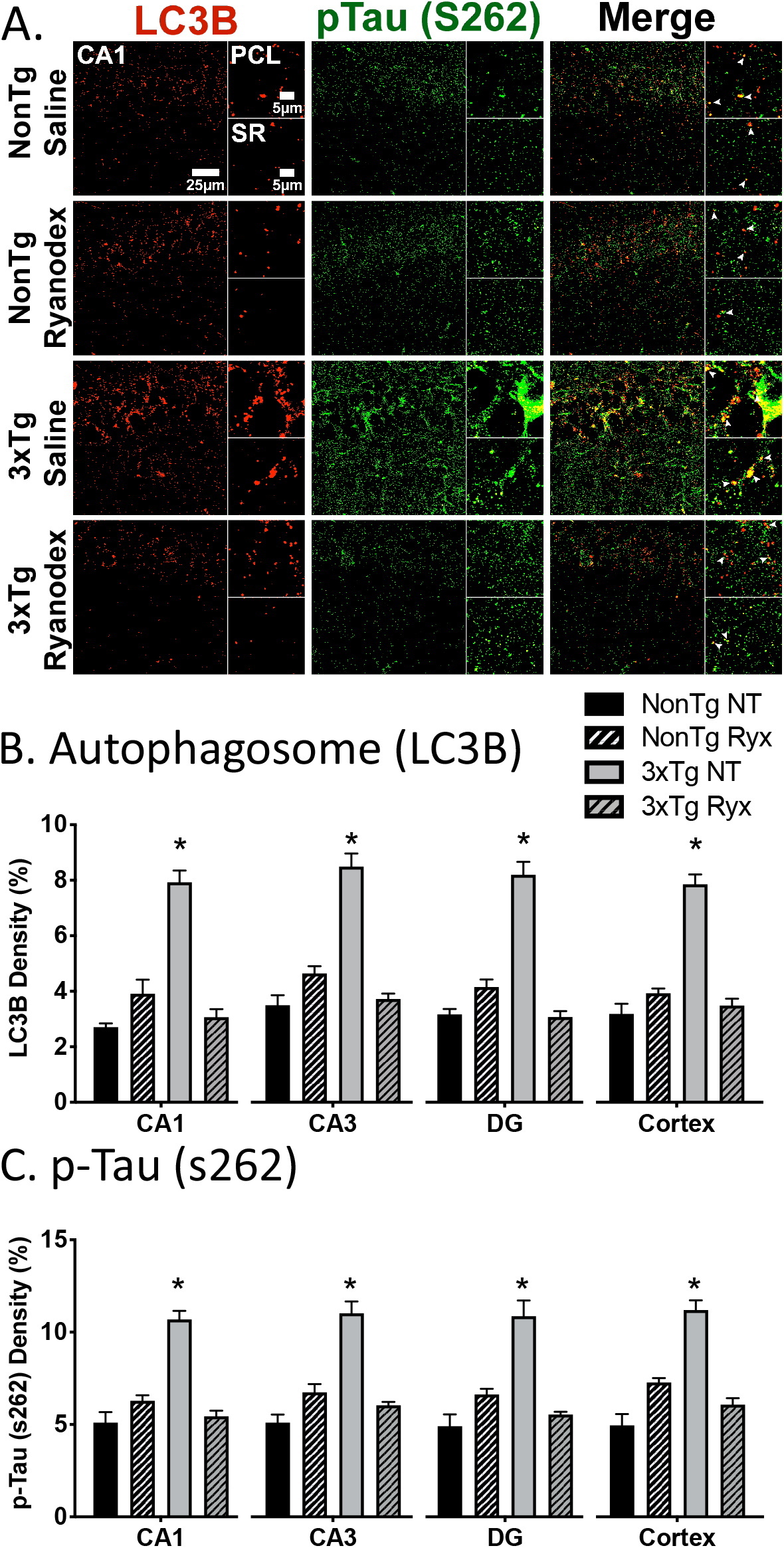
Increased autophagosomes in AD mice are restored after normalizing Ca^2+^ levels. Autophagic clearance of hyperphosphorylated tau is dependent on regulated RyR-Ca^2+^ signaling. Hippocampus from 3-4-month-old 3xTg-AD mice and age-matched NTg controls were immunolabled for autophagosomes (LC3B) with early stage phosphorylated tau (S262). (A) Representative high-resolution images were taken on Leica SP8 63X with 441X zoom of the primary cellular layer (PCL) and stratum radiatum (SR) of the CA1 field. Quantification showed decreased fluorescence density (% of threshold area over background) for (B) LC3B-II and (C) p-tau S262 in multiple regions of the hippocampus (CA1, CA3, DG) and cortex; which was rescued with Ryanodex treatment (10mg/kg; 30-day ip). (NTg Vehicle, black bar, n=8 animals; NTg Ryanodex, black pattern bar, n=8 animals; 3xTg Vehicle, grey bar, n=8 animals; 3xTg Ryanodex, grey pattern bar, n=8 animals) *\*p <0.05*.

Autophagosomes are responsible for engulfing abnormal, pathogenic proteins, such as intracellular hyperphosphorylated tau (p-tau), and thus, accumulation of abnormal proteins would be an expected consequence of impaired proteolysis. To explore this, immunolabeling of early p-tau (S262 antibody) was conducted and increased staining was observed in the dorsal hippocampus and cortex of 3xTg-AD mice, effects that were normalized with Ryanodex treatment in 3xTg-AD mice comparable to NTg levels **(Figure 6A,C;** n=8, CA1 (F_(3,28)_=37.08, p<0.0001); CA3 (F(3, 28)=33.16, p<0.0001); DG (F_(3,28)_=22.82, p<0.0001); Cortex (F_(3,28)_=35.47, p<0.0001). To determine if these deficits translate to human pathology, autophagosome levels were measured in the HiN using the live-cell DAP autophagy sensor. Significant aggregation of autophagosomes, as measured by DAP-green staining density, was observed in AD HiN as compared to non-AD HiN. Normalizing RyR-Ca^2+^ release with Ryanodex rescued these deficits in AD HiN to levels comparable to non-AD HiN **(Figure 7A,C;** n=5, (F_(3,15)_=7.32). Collectively this demonstrates that aberrant RyR-Ca^2+^ release disrupts autophagy and results in a build-up of autophagosomes containing aberrant proteins in AD murine and human neurons, which is resolved with normalizing intracellular Ca^2+^ levels.

**Figure 7.**
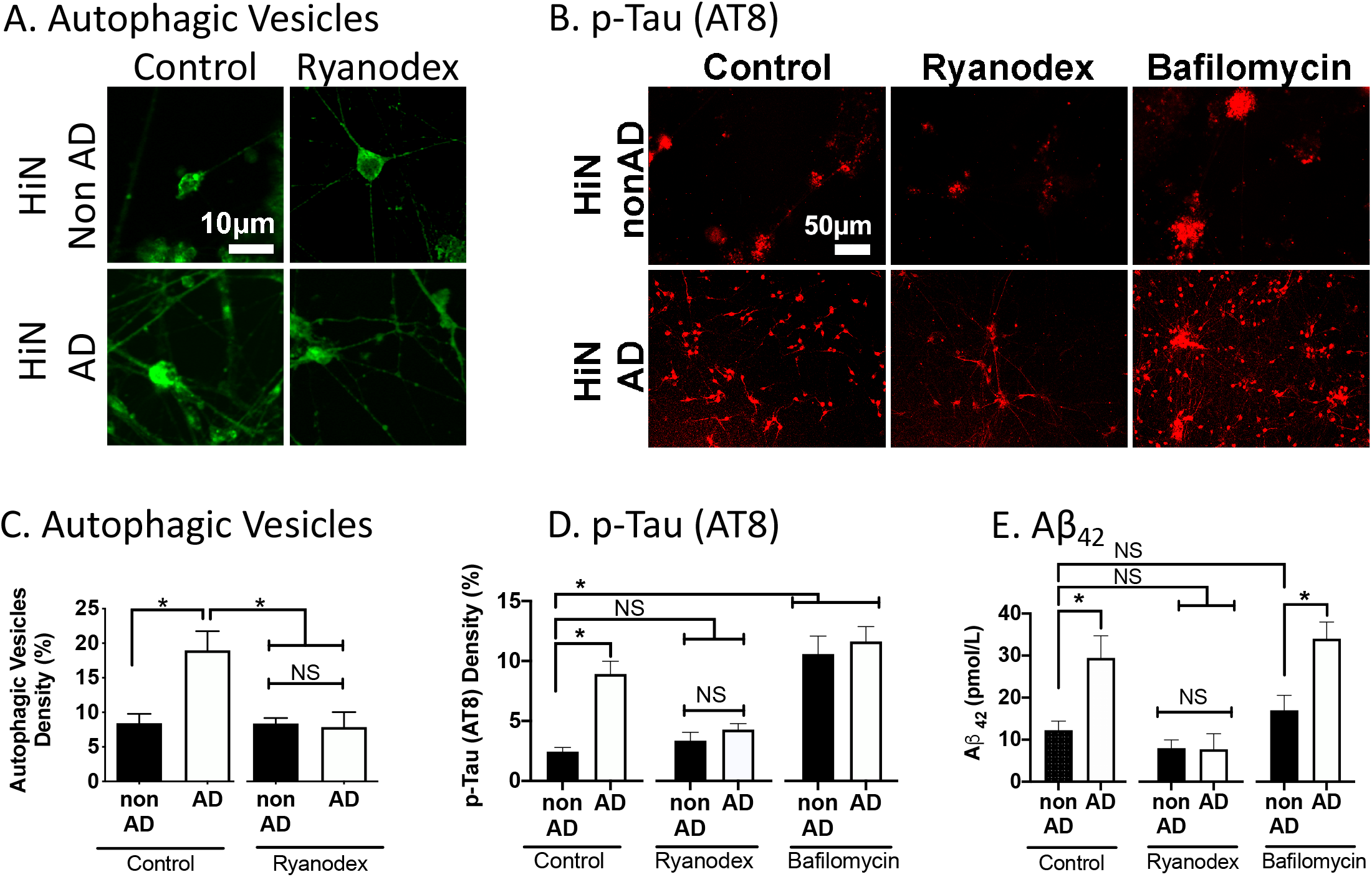
Autophagy inhibition exacerbates hyperphosphorylated tau aggregation in human neurons. Regulated RyR-Ca^2+^ release is necessary for autophagy-mediated clearance of intracellular protein aggregates. (A) Representative images of autophagic vesicles, measured by DAP-green autophagy detection dye in non-AD and AD HiN. (C) Quantification of fluorescence density of autophagic vesicles (% threshold area over background) show increased density of autophagic vesicles in AD HiN, which are rescued with Ryanodex treatment (n=5 coverslips/treatment). (B) Representative images of fixed HiN cultures immunolabeled for hyperphosphorylated tau (AT8). (D) Quantification of fluorescence density of hyperphosphorylated tau (% threshold area over background) show increased AT8 in AD HiN. Inhibition of vATPase with bafilomycin (125nM) exacerbated p-tau in non-AD HiN. Ryanodex (10μM) treatment reduced the elevated AD-associated hyperphosphorylated tau in AD HiN. (n=10 coverslips/treatment). (C) Aβ_42_ specific ELISA of the supernatant from the non-AD and AD HiN showed increased secreted Aβ_42_ in AD neurons. Bafilomycin treatment did not exacerbate secreted Aβ_42_ levels in non-AD HiN, showing that blocking autophagy has a more profound effect on intracellular deposits than extracellular deposits. Additionally, Ryanodex (10μM) treatment significantly reduced extracellular Aβ_42_ levels in AD HiN comparable to non-AD HiN (n=8 wells/treatment). *\*p <0.05.*

### Inhibiting autophagic clearance increases AD pathogenic species in human AD models

Pathogenic protein fragments accumulate as a result of defective autophagy and disrupted lysosomal function. To test this directly, we measured AT8-immunolabeled p-tau in the HiN. Phosphorylated tau (AT8) was increased in AD HiN as compared to non-AD HiN and Ryanodex reduced p-tau expression to non-AD levels. Interestingly, vATPase inhibition with bafilomycin elevated p-tau levels in non-AD HiN to levels similar to AD HiN, indicating that lysosomal alkalization and autophagic disruption exacerbate intracellular protein aggregates (**Figure 7B,D;** n=10, F_(5,54)_=16.58, p<0.0001). To measure extracellular AD protein aggregates, an ELISA was performed for extracellular Aβ_42_ species on supernatants collected from non-AD and AD HiN. Increased levels of secreted Aβ_42_ were seen in AD HiN as compared to non-AD HiN samples. Normalizing intracellular Ca^2+^ levels with Ryanodex significantly decreased extracellular Aβ_42_ levels. However, blocking vATPase with bafilomycin did not significantly alter secreted Aβ_42_ levels in non-AD nor AD HiN; suggesting that lysosomal defects have a profound effect on intracellular protein aggregates **(Figure 7E;** n=8, F_(5,49)_=9.28, p<0.0001). Together, this study demonstrates that aberrant RyR-Ca^2+^ release underlies lysosomal dysfunction and halts autophagic-mediated clearance of intracellular pathogenic proteins, furthering AD proteinopathy.

## Discussion

In this study, we reveal a novel mechanism through which defective lysosomal acidification contributes to pathological accumulation of protein fragments in a Ca^2+^-dependent manner in AD. Furthermore, pharmacological suppression of RyR-Ca^2+^ release rescued lysosomal acidity, proteolytic function and autophagic clearance. Through validation of these pathogenic mechanisms in HiN from AD patients, novel targets for therapeutic intervention are revealed **(Figure 8)**.

**Figure 8.**
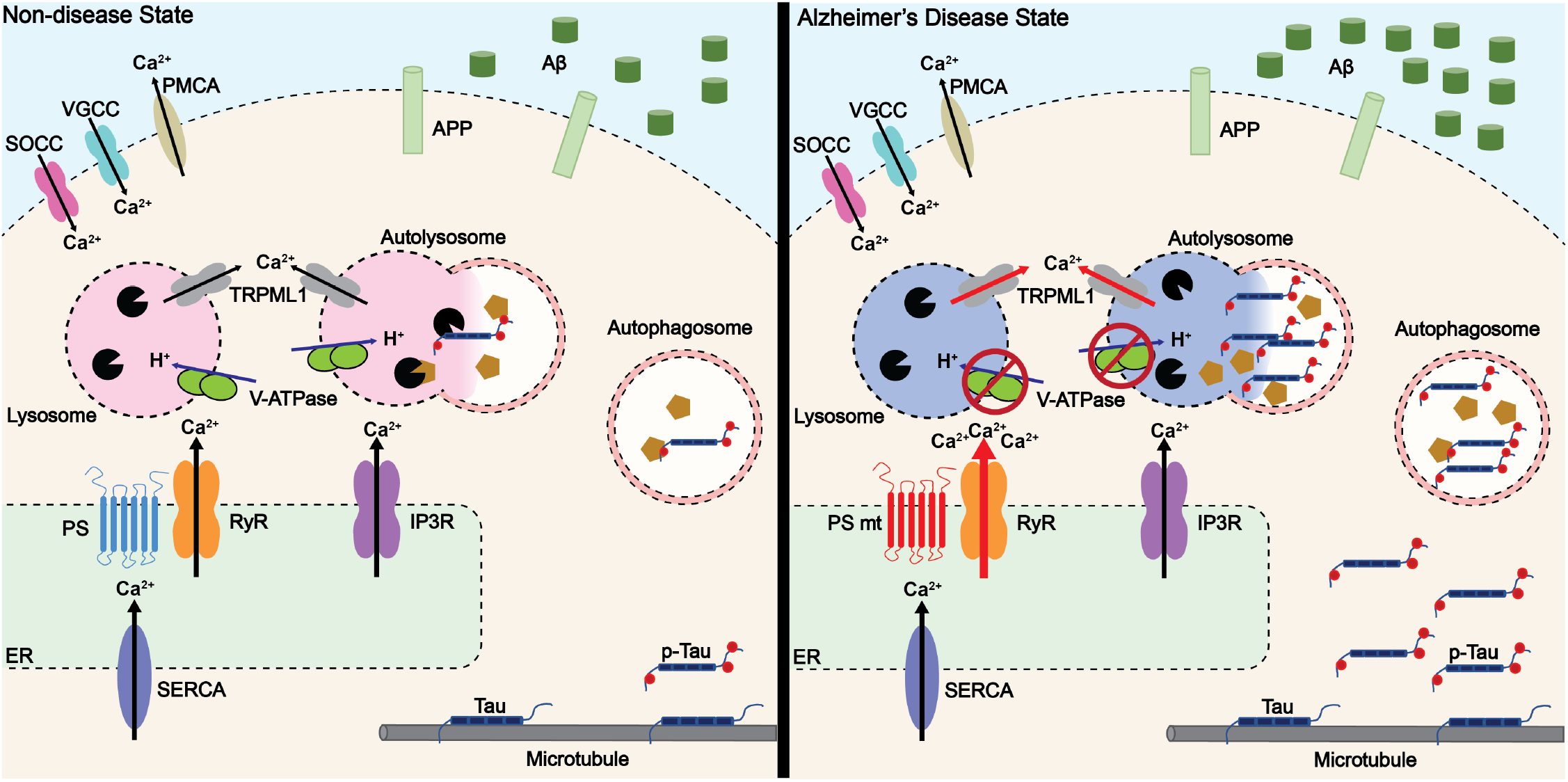
Aberrant RyR-Ca^2+^ disrupts lysosomal-autophagic mediated degradation of intracellular protein aggregates in AD. Protein handling via lysosomal-autophagosome pathway is disrupted in early stages of Alzheimer’s disease. Dysregulated RyR-Ca^2+^ signaling leads to elevated cytosolic Ca^2+^ concentration. This rise of Ca^2+^ disrupts lysosomal vATPase proton pump, resulting in alkalization of the lysosomal lumen. The alkaline environment is not conducive for lysosomal protease, cathepsin, hydrolase activity thereby disrupting proteolysis of cellular debris. Autophagic clearance is halted and turnover of bulky, misfolded proteins and damaged organelles is disrupted. Intracellular aggregates such as hyperphosphorylated tau accumulate within autophagic vacuoles furthering AD proteinopathy.

### Defects in lysosomal acidification

Sufficiently acidified lysosomes are essential for numerous cellular functions including degradation and recycling of intracellular debris and waste products, synaptic plasticity and maintaining sufficient Ca^2+^ reservoirs (24, 56, 57). Acidification is maintained by the vATPase proton pump, and previous reports show that reduced V0a1 subunit expression in mutant PS1-expressing cells results in alkalized lysosomes and blunted autophagy (16, 17, 23, 25). Coinciding with these findings, our study demonstrates decreased lysosomal vATPase levels in 3xTg-AD mouse brains prior to detectable histopathology (**Figure 1)**, and, of relevance to human pathophysiological mechanisms, similar findings were found in HiN from AD patients with PS1 mutations (**Figure 4)**.

A particularly novel finding that normalizing excess AD-associated RyR-evoked Ca^2+^ release restored vATPase levels in 3xTg-AD mice and AD HiN to that of the NTg and non-AD HiN controls. This directly implicates intracellular Ca^2+^ dysregulation in altering lysosomal vATPase expression levels, and reinforces the important role of inter-organelle Ca^2+^ communication for vATPase trafficking to lysosomes, assembly of both domains, or protein biogenesis. With the aid of PS1, post-translational modifications of proteins, such as *N*-glycosylation, is required for maturation and trafficking of V0a1 to the lysosomes (16). Additionally, recent biochemical studies reveal multiple glycosylation sites within the structure of human vATPase, especially in the subunit V0a subunit, which are critical for assembly, stability, protein folding during vATPase biogenesis, and protection from proteolytic activity within lysosomes (58, 59). As this vATPase N-glycosylation occurs in the ER and Golgi apparatus, it is likely that ER-Ca^2+^ signaling has a key regulatory role for post translational modifications that affect its assembly and insertion to the lysosome.

In AD neurons, the reduced capacity of lysosomes to import protons impairs pH regulation resulting in relatively alkaline baseline pH levels compared to control neurons. Notably, the abnormal pH was restored to acidic levels when RyR-Ca^2+^ signaling was normalized. In control neurons **(Figure 3,4)** and RyR2/3 overexpressing cell models **(Figure 2)**, lysosome pH was alkalized after evoked RyR-Ca^2+^ release, which was again attenuated with normalization of RyR-Ca^2+^ release, revealing a new component of ER and lysosome crosstalk that plays a critical role in protein handling. The close temporal coupling between ER-Ca^2+^ release and the subsequent lysosome pH response is demonstrated in the overlay traces **(Figure 2)**, and demonstrates that lysosome alkalization occurred immediately after evoking RyR Ca^2+^ release, speaking to the close proximity and direct coupling of the ER to lysosomes. In polarized neurons, the contact sites between lysosomal and ER Ca^2+^ efflux channels form Ca^2+^ microdomains that play a crucial role in distribution and trafficking of late endosomes/lysosomes, lipid transfer, and autophagy regulation (10, 38–40, 60, 61). Disrupting this Ca^2+^ microdomain homeostasis, such as through excessive RyR-evoked Ca^2+^ release as described in AD, thus directly interferes with lysosomal ionic flux and pH regulation.

The excess RyR-Ca^2+^ release in AD likely disrupts vATPase function through interference with ion cotransporters and exchangers, causing an imbalance in local ionic homeostasis, thus reducing the driving force of H^+^ influx via vATPase. The critical role of ion gradients in lysosome function is also demonstrated in a recent study that implicates the chloride channel, CLC7, as a counterion conductance channel that regulates lysosomal pH, and loss of this channel exacerbates lysosomal dysfunction (62). The balance of ion channel regulation thus plays a central role in maintaining proteolysis capacity and clearance of undesirable cellular fragments, and when ionic homeostasis is disrupted, aberrant protein aggregates accumulate, such as seen in neurodegenerative disorders such as Alzheimer’s disease.

As Ca^2+^ harboring organelles, lysosome Ca^2+^ stores ([~0.5mM]) are utilized for localized Ca^2+^ signaling, fusion to cargo vesicles, endocytosis, and lysosomal biogenesis (10). Nicotinic acid adenine dinucleotide phosphate (NAADP)-sensitive channels, such as two-pore calcium channel protein (TPC) and TRPML serve as dominant Ca^2+^ efflux channels (63, 64). Ca^2+^/H^+^ exchangers or Ca^2+^ transporters are thought to establish the lysosomal Ca^2+^ and H^+^ gradient and balance vATPase function (65, 66). In this scenario, the high density of tightly coupled ion channels regulating lysosome function generates a sensitive dynamic in that alterations in any one ion class can have rather profound effects on ionic homeostasis and overall organelle function. Additionally, the vATPase is structurally complex and many of the Ca^2+^ regulated domains and interaction partners are still under investigation. Thus, while the exact mechanism by which RyR-Ca^2+^ affects lysosomal pH is under investigation, direct and indirect effects on vATPase structure and function are likely mechanisms.

The cellular localization of the lysosome also informs about its degradative status, with mature lysosomes in the soma being the most acidic (pH 4.5-5) with the greatest catabolic activity. In distal processes, autophagosome and endosomes are “immature lysosomes” that progressively become more acidic in their retrograde transport, reveling a gradation of acidification (67–70). In this study, lysosomal pH fluctuations were primarily measured from mature lysosomes within the soma, and thus, in AD neurons, the most degradative lysosome population is rendered ineffective with insufficient proteolysis capacity elsewhere in the cell. Furthermore, as excessive ER Ca^2+^ release is present across multiple neuronal compartments in AD models, including soma, dendrites, spines, and axon terminals, this widespread Ca^2+^ dyshomeostasis can inflict impairments in the trafficking of these cargo vesicles, further exacerbating proteinopathy cascades. Correspondingly, our findings reveal accumulated autophagosomes within the dendrites in the CA1 stratum radiatum hippocampal subfield in 3xTg-AD mice **(Figure 6)**.

### Defects in autophagosomal-lysosomal mediated degradation

Lysosomal protease activity is disrupted in AD human neurons (**Figure 5)**, and aligns with studies demonstrating inactivation of lysosomal hydrolytic enzymes, including cathepsin D,B,L, in neurodegenerative diseases (71, 72). The hindered proteolytic activity results in profuse accumulation of autophagic vesicles containing aberrant protein deposits, like p-tau, in AD mouse models and HiN from AD patients (**Figure 6–7**; (73, 74)). Here we expanded upon these mechanisms to demonstrate the contribution of aberrant Ca^2+^ signaling to protein mishandling in AD pathogenesis by rescuing autophagy and clearing p-tau by normalizing RyR-Ca^2+^ signaling **(Figure 6–7)**. These findings coincide with a recent study demonstrating that RyR-Ca^2+^ activity suppressed autophagic flux and resulted in accumulation of autophagic vesicles (75). Inhibiting autophagy with bafilomycin exacerbated hyperphosphorylated-tau accumulation in HiN; interestingly however, there were no effects on secreted Aβ_42_ levels **(Figure 7)**. Notably, the APP holoprotein is a type 1 transmembrane protein with the Aβ cleavage peptide spanning the membrane and the extracellular space; thus, secreted amyloid peptides are likely less accessible to cytosolic autophagosome-mediated clearance.

However, a recent study showed that accumulation of intracellular Aβ/APP-βCTF within de-acidified autolysosomes occurs in early stages of AD, and the impaired clearance results in large autophagic vesicle packs that contribute to senile plaque formation (76). Additional groups have shown that the cytosolic APP fragments (APP intracellular domain, AICD), a result of γ-secretase activity on the C99 fragment of APP, accumulate within acidic compartments such as lysosomes and endosomes in neurons (77). This suggests that localized large ER Ca^2+^ bursts from RyR can have compounding repercussions on autophagic clearance of intracellular protein aggregates in AD neurons. Of note, normalization of RyR-Ca^2+^ signaling restored levels of secreted Aβ_42_ levels comparable to non-AD levels, displaying a lysosomal independent mechanism for secreted Aβ_42_ regulation. Here, the mechanism may reflect the potentiation of APP phosphorylation through Ca^2+^-mediated positive regulation of β- and γ-secretase activities, and thus normalizing Ca^2+^ levels will result in the lowering of C99 and pathological Aβ production (36, 78).

Gene-level analysis also reveals alterations in protein handling pathways in AD populations, which may serve as a biomarker to identify at-risk individuals, or as possible targets for gene therapy. Transcriptomic profiles from biobanks including the Rush Memory and Aging Project (MAP), Religions Orders Study (ROS), and Mount Sinai Brain Bank (MSBB) reveal clusters of genes that associate with pathological protein handling (79–82). Specifically, altered expression of BIN1, ABCA7, and SORL1 is linked to blunted neuronal endosomal trafficking, reduced degradative potential of lysosomes, and decreased autophagy, thus furthering AD proteinopathy (83). Importantly, in AD populations, the vATPase subunit V1A, and transcription factor EB (TFEB), a master regulator for lysosomal biogenesis, were downregulated (81, 82).

Here, we propose a novel mechanism by which dysregulated RyR-mediated Ca^2+^ release drives proteinopathy in AD. ER-Ca^2+^ dyshomeostasis is described in a variety of AD animal, cellular models, and human neurons prior to plaque and tangle formation (27, 44, 84–87), and is caused in part through post-translational modifications such as oxidation, nitrosylation, and phosphorylation (32, 53, 88–91). This is evident in the AD brain where mitochondrial-mediated oxidation of the RyR (92), and PKA phosphorylation (53) contribute to exaggerated RyR-Ca^2+^ signaling. Thus, by preventing aberrant Ca^2+^ signaling in AD neurons, a broad array of therapeutic outcomes emerge.

## Materials and Methods

### Mouse Models and Ryanodex Treatments

Three -to four-month-old male and female 3xTg-AD [APP_swe_, Tau_P301L_, and PS1_M146V_KI; (41)] mice and age-matched non-transgenic control mice [C57BL6/J9] were used in this study. Mice were housed at the RFUMS Biological Resource Facility in accordance with IACUC regulations, and kept on a 12:12h light/dark cycle with food and water available ad libitum. A nanocrystal formulation of dantrolene, Ryanodex (Lyotropic Therapeutics Inc.) was administered intraperitoneally at 10mg/kg to mice for four weeks starting at three months of age. Vehicle-control 3xTg-AD and NTg mice were administered 0.1mM sterile saline. Body weight was taken every five days during treatment to monitor for significant health issues. There were no differences in body weight during the course of the study, or in brain weight or brain:body weight in treated animals p<0.05 (35).

### Cell Culture

Stable, inducible HEK293T cells expressing WT RyR2 or RyR3 were obtained from Dr. Wayne Chen and are well-characterized (42, 43). Cells were grown and maintained in Dulbecco’s modified Eagle’s medium (DMEM) supplemented with 0.1 mM minimum Eagle’s medium nonessential amino acids, 4 mM L-glutamine, 100 IU/ml penicillin 100/mg of streptomycin, 4.5 g glucose/liter, and 10% fetal calf serum, at 37°C under 5% CO2. Cells expressing the various RyR isoforms were selected using 200μg/ml hygromycin. Cells were plated on glass coverslips coated with 0.1% Poly-L-lysine or 96-well Corning black, clear bottom plates and given doxycycline to induce RyR expression 48hrs prior to use. Immunofluorescence was used to confirm the presence of RyR2 and RyR3 (data not shown).

### Primary Neuronal Culture

Hippocampal neurons were obtained from post-natal day 1 or 2 mouse pups. Hippocampi were dissected and washed with ice-cold Hanks’ balanced salt solution (HBSS) without Mg^2+^ or Ca^2+^ and supplemented with 10 mM HEPES at pH 7.3. Following a 30-min incubation with 0.25% trypsin at 37°C, the hippocampi were again washed with the warm HBSS + 10mM HEPES solution and dissociated. They were seeded on Matrigel treated coverslips at 50,000 cells per well. Cells were grown in seeding medium (neurobasal medium containing 10% FBS, 100 IU/ml penicillin, 100 mg/ml streptomycin, 2mM glutamax and 1:50 B27 supplement (Life Technologies). After 10-12 days in culture, cells were incubated with indicated concentrations of vehicle, Ryanodex, or bafilomycin A1 before running physiological experiments. Neuronal cultures were stained for NeuN to confirm neuronal identity.

### iPSC and HiN Generation

Generation of iPSC-derived HiN has been previously described (44). Briefly, normal control- and familial-AD patient-fibroblasts (A246E PS1 mutation) obtained from the Coriell Institute were converted to iPSCs using gene delivery of the Yamanaka factors by vector or RNA transfection (ReproRNA-OKSGM kit). Fibroblasts were transfected and cultured for 14–21 days in REPRO-TeSR medium. Viable iPSC colonies were converted to HiN by lentiviral vectors containing the transcription factor NGN2. HiNs were selected for lentiviral transduction by puromycin resistance, allowed to mature over 14–28 days in Neural Basal Media (ThermoFisher, Waltham, MA, USA) with doxycycline, brain-derived neurotrophic factor (BDNF) and neurotrophin 3 (NT3) (PeproTech, Rocky Hill, NJ, USA). Induced neurons were fed with 50:50 mixture of astrocyte conditioned medium (provided by Dr. Allison Ebert). HiN were grown on matrigel coated coverslips or 24-well (Costar) plate.

### Lysosomal pH and Autophagic Vesicle Imaging

Lysosomal pH was measured using a dextran-conjugated ratiometric Lysosensor dye targeted to lysosomes, optimized for lysosomal pH range, and sensitive to minute changes in lysosomal pH (9, 16). 0.05 mg/ml Lysosensor Yellow/Blue-dextran (Invitrogen) was added and incubated for 4 hours at 37°C with 5% CO2. Cells were washed with HBSS+ 10mM HEPES buffer and then read either on FlexStation 3 (Molecular Devices) or an inverted Nikon Eclipse Ti2 microscope coupled to a mercury light source (Nikon Intensilight C-HGFIE) and Lambda 10-B Sutter filter switch. Excitation wavelength was 355nm; emission ratio at 440nm/535nm. The standard curve to measure pH values was generated using a modified protocol of the intracellular pH Kit (Life Technologies) in which cells were treated for 15 min with 10μM Valinomycin and 10μM Nigericin, in either the given buffers or MES buffer (5mM NaCl, 115mM KCL, 1.3mM MgSO4, 25mM MES) adjusted to pH 3.5 with HCl (9). Calibration measurements are made simultaneously with pH measurements in adjacent wells, controlling for time or dye-dependent effects. To confirm lysosome identity (16, 45), neurons were incubated with LysoTracker Red DND-99 (1μM; 2hrs; ThermoFisher) then fixed with 4% PFA and imaged with a 20X objective. Images were analyzed on MetamMorph software (version 7) and presented as mean fluorescent density above background.

To measure autophagic vesicles, neurons were incubated with DAPGreen-Autophagy Detection dye (0.1μM; 6 hrs.; Dojindo Molecular Technologies, Inc). Neurons were then imaged with a 20X air objective using an inverted Nikon Eclipse Ti2 microscope as described above.

### Calcium Imaging

Procedures were performed as previously described (44). Briefly, cells were incubated with Fura-2AM (5μM; 1hr; Invitrogen) then imaged on a modified upright Olympus BX51WI microscope. Cells were continuously perfused with oxygenated (95% O_2_-5% CO_2_) artificial cerebrospinal fluid (aCSF) [125mM NaCl, 2.5 KCl, 1.25mM KH_2_PO_4_, 10mM dextrose, 25mM NaHCO_3_, 2mM CaCl_2_, and 1.2mM MgSO_4_, pH 7.4] at room temperature. The microscope is coupled to a xenon light source (Excelitas Technologies) and DG5 Sutter filter switch. The fluorescence responses were captured using a 40X water-immersion Olympus objective and analyzed using Nikon NIS-Elements Software (AR package). Bath application of caffeine (10mM, Sigma) was used to evoke RyR-evoked calcium release. Ryanodex (10μM; Lyotropic Therapeutics Inc.) was dissolved in aCSF and used as a negative allosteric modulator of RyR. Fura-2AM calcium responses are reported as peak 340/380 ratio over baseline response after background subtraction.

### Immunohistochemistry

Mice were anesthetized with urethane (20mg/kg) and then transcardially perfused with ice-cold phosphate-buffered saline and then 4% paraformaldehyde. Brains were removed and post-fixed for 24 hours and replaced with 30% sucrose solution for 3 days. Brains were cut into coronal sections on a Leica SM 2000R microtome with a freezing stage at 40μM thickness. Free-floating hippocampal sections were permeabilized with 1X TBS 1.0% Triton-X-100 (15min; room temperature) then blocked with 10% goat serum (1hr; room temperature) on a platform rocker, then incubated in primary antibody [V0a1 or V1B2 (1:500; custom-generated antibodies developed and validated by Drs. Beaman and Gilman-Sachs (46, 47); LC3B (1:500; Abcam 48394)] overnight at 4°C on a platform rocker. Sections were washed with 1X TBS 0.1% Triton-X and incubated in secondary antibody [Alexa Fluor 555 (1:1000; Invitrogen A21428)] for 1hr at room temperature on a platform rocker. Sections were rinsed with 1X TBS 0.1% Triton-X and incubated in the second primary antibody [phospho-Tau S262 (1:500; Invitrogen 44-750G) Lamp1 (1:500; Cell Signaling CS4H11)] diluted in 1X TBS 0.1% Triton-X + 1% goat serum for overnight at 4°C. Sections were rinsed 1X TBS 0.1% Triton-X and incubated in the second secondary antibody [Alexa Fluor 488 (1:1000; Invitrogen A-11008)] diluted in 1X TBS 0.1% Triton-X + 1% goat serum for 1 hour at room temperature. Sections were rinsed in 1X TBS and stained with 1:10000 DAPI diluted in 1xTBS for 5 minutes. Sections were washed in 1X TBS and mounted with PVA-DABCO for microscopy. High resolution confocal images were obtained using a 63x oil objective lens on Leica SP8 imaging system. MetaMorph software (v. 7) was used to quantify the staining density above background threshold and percent overlap for colocalization studies.

### Immunocytochemistry

Procedures were performed as previously described (44). Briefly, HiNs were fixed overnight in 4% paraformaldehyde, permeabilized with 1X TBS 1% Triton-X-100 (5min; room temperature) then blocked using 10% normal goat serum (1hr; room temperature) on a platform rocker. HiNs were incubated with primary antibody (hyperphosphorylated tau: AT8 (1:1000; ThermoFisher MN1020) overnight at 4 °C on a platform rocker. HiN were then rinsed with 1X TBS and incubated in secondary antibody [Alexafluor 594 (1:1000 ThermoFisher)] at room temperature on a platform rocker for 1hr. Fluorescent images were then collected using a 20X air objective using an inverted Nikon Eclipse Ti2 microscope coupled to a mercury light source (Nikon Intensilight C-HGFIE), captured with Nikon Elements software (AR package). Images was analyzed on MetamMorph software (v.7) and presented as mean fluorescent density above background.

### Enzyme Linked Immunosorbent Assays

Enzyme linked immunabsorbent assays (ELISA) were performed on supernatants from mature HiN cultures using a capture antibody kit specific for Aβ42 (WAKO Chemicals) as per manufacturer’s instructions. Detection of Aβ42 was run on FlexStation3 (Molecular Devices) and determined by colorimetric change as compared to standard curve generation.

### Experimental Design and Statistical analyses

Statistical analysis was performed using Graph Pad Prism 7. Data are represented as mean ± SE. Unpaired T-tests, One-way ANOVA or two-way ANOVA, with Tukey’s post hoc analysis, were performed where appropriate. Statistical significance was set at p<0.05.

## Acknowledgments

We would like to thank Dr. Wayne Chen, University of Calgary, for graciously providing the HEK293T RyR cell lines, Dr. Allison Ebert, Medical College of Wisconsin, for graciously supplying the astrocyte cultured medium, and Dr. Kenneth Beaman, Rosalind Franklin University of Medicine and Science for graciously generating the custom vATPase antibodies, all of which were used in immunoassays, optical imaging, and production of human neurons.

